# SwarmTCR: a computational approach to predict the specificity of T Cell Receptors

**DOI:** 10.1101/2020.11.05.370312

**Authors:** Ryan Ehrlich, Larisa Kamga, Anna Gil, Katherine Luzuriaga, Liisa Selin, Dario Ghersi

## Abstract

**Motivation:** Computationally predicting the specificity of T cell receptors can be a powerful tool to shed light on the immune response against infectious diseases and cancers, autoimmunity, cancer immunotherapy, and immunopathology. With more T cell receptor sequence data becoming available, the need for bioinformatics approaches to tackle this problem is even more pressing. Here we present SwarmTCR, a method that uses labeled sequence data to predict the specificity of T cell receptors using a nearest-neighbor approach. SwarmTCR works by optimizing the weights of the individual CDR regions to maximize classification performance.

**Results:** We compared the performance of SwarmTCR against a state-of-the-art method (TCRdist) and showed that SwarmTCR performed significantly better on epitopes EBV-BRLF1_300_, EBV-BRLF1_109_, NS4B_214–222_ with single cell data and epitopes EBV-BRLF1_300_, EBV-BRLF1_109_, IAV-M1_58_ with bulk sequencing data (α and β chains). In addition, we show that the weights returned by SwarmTCR are biologically interpretable.

**Availability:** SwarmTCR is distributed freely under the terms of the GPL-3 license. The source code and all sequencing data are available at GitHub (https://github.com/thecodingdoc/SwarmTCR)

## Introduction

The adaptive immune system plays a critical role in curbing infections and in cancer immunosurveillance, defined as the patrolling of the body by the immune system with the active elimination of precancerous and cancerous cell.^1^ CD8+ T lymphocytes are one of the key cell types involved in antiviral responses and cancer immunosurveillance. They perform their function by binding to small peptides presented on the surface of highly polymorphic molecules known as the Major Histocompatibility Complex (MHC) using T Cell Receptors (TCR). TCRs are transmembrane proteins contain either α and β or γ and δ chains, within which are three loops called Complementarity Determining Regions (CDRs). CDR loops are characterized by both germline loops (CDR1 and CDR2) and the hyper-variable CDR3 loop, which is the product of somatic recombination.^2,3^ These CDR loops are responsible for interacting with the peptide/MHC (pMHC) complex. The diversity of TCR sequences is mostly focused on the CDR regions and is very large, with numbers in human that are thought to exceed 10^20^ possible distinct receptors.^4^

The collection of TCRs possessed by an individual is known as the T cell repertoire, which is shaped over time by the history of infections in combination with stochastic factors, and is in turn responsible for determining the outcome of an immune response. With access to enough sequencing data, it is now theoretically possible to predict the specificity of the T cell repertoire of an individual using sequence information alone.^2^ In other words, we could determine the identity of the peptide(s) that each TCR is capable of recognizing by computationally analyzing the TCR sequences from an individual. Two main technologies are currently available for sequencing TCRs: (1) single cell (SC) sequencing; and (2) bulk sequencing (BS). SC TCR sequencing technology allows to reconstruct the complete sequence of a TCR with paired α and β chain sequence information, but its cost is still limiting the amount of available data. In contrast, BS technology is more affordable and has yielded substantially larger amounts of data. However, reconstructing the correct α and β chain pairs within a TCR is not possible with this technology.

Being able to map the specificity of human repertoires can equip us with powerful new tools for studying autoimmunity, cancer immunotherapy, and immunopathology.^5^ However, for these methods to be broadly applicable it is critical to sample T cell repertoires deeply and in multiple individuals, as well as to account for the diversity of binding topologies to pMHCs with computational approaches. Here we introduce SwarmTCR, a computational method to predict the specificity of TCRs for class I MHC/peptide complexes that outperforms the state-of-the-art approach TCRdist^2^ on both SC and BS data.

TCRdist uses a nearest-neighbor approach with a pairwise sequence alignment score between TCRs as a proximity measure. The two chains are weighted equally, and the CDR3 region is weighted three times more than the other CDR regions. While this is a reasonable choice considering the importance of CDR3 for peptide binding, it does not take into consideration the fact that the two chains and the regions within them might have different levels of involvement in binding to the pMHC, depending upon the peptide being presented and the MHC type. In a recent study, we curated a non-redundant set of TCR/pMHC crystal structures and explored binding topologies of TCR/pMHC complexes and the number of contact residues (≤ 4.5 Å^6^) made by α and β chains with pMHC.^7^ Our results indicated a wide range of TCR binding angles and a variable use of the α (7 - 25 contacts) and β (6 - 22 contacts) chains in making contacts with the pMHC. We also computed the number of alpha and beta contacts to the pMHC, determining a ratio of contacts (α/β ratio) for each structure. In some complexes, the α chain had a much larger number of interactions with the pMHC than the β chain, whereas in other complexes the β made more interactions than the α chain. In other complexes, we saw an almost equal number of α /β interactions with the pMHC ~15 contacts per chain. Taken together, these results suggest a wide range of binding recognition modes, which should be reflected in a computational method to predict TCR binding specificities.

As a first step to leverage these findings, we developed SwarmTCR, a method to predict TCR specificity that automatically learns the optimal set of weights to assign to each CDR region based on classification accuracy in a cross-validation setting (Figure 1). In addition to CDR1, CDR2, and CDR3, the method also incorporates the CDR2.5 region (a loop between CDR2 and CDR3 that can interact with the pMHC, as discussed in TCRdist^2^) for a total of four weights per chain. By directly optimizing the weights for CDR regions in a peptide-specific fashion, our method automatically accounts for the diversity in pMHC recognition that is documented in crystal structures (see Methods).

**Figure 1.**
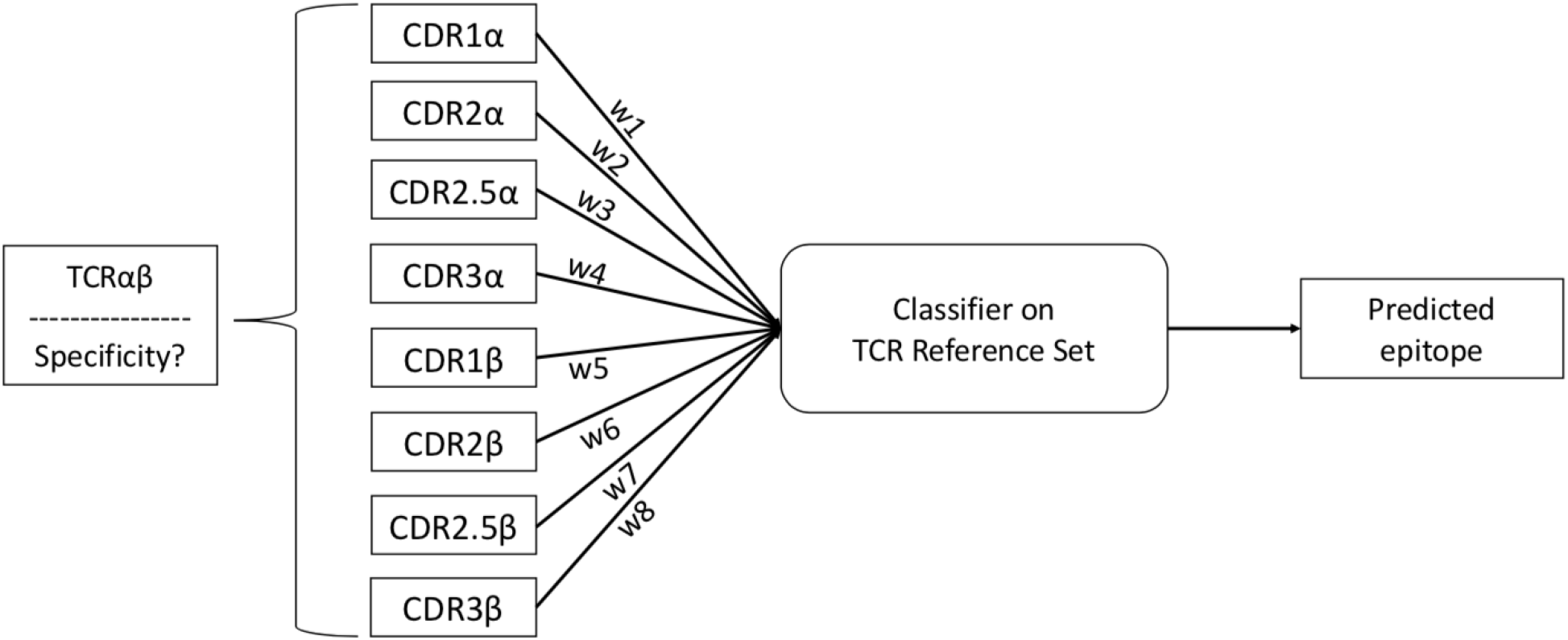
Here the SC model is illustrated. Weights for each CDR loop are determined by the optimization step and subsequently tested on the testing set to assess peptide prediction performance.

We applied our method to SC and BS data and compared its performance against that of TCRdist. In addition to performing consistently better than TCRdist, the weights returned by SwarmTCR in SC sequencing data can potentially inform the user about the contribution of the two chains in recognizing the pMHC complex.

## Methods

### TCRs sequence data

CD8+ TCR SC and BS data were collected from: (1) the Selin and Luzuriaga Labs at UMASS; (2) VDJdb^8^ and (3) IEDB^9^.

Data acquired from the Selin and Luzuriaga labs contained TCRs isolated from HLA A0201-restricted, naïve and peptide-specific CD8+ T cells binding to YVL (EBV-BRLF1_109_: HLA-A0201 restricted, peptide: YVLDHLIVV), GLC (EBV-BRLF1_300_:HLA-AO2O1 restricted, peptide: GLCTLVAML), and GIL (IAV-M1_58_: HLA-A0201 restricted, peptide: GILGFVFTL).

Human data from VDJdb was downloaded in January 2018, where paired TCR information is denoted by matching index values and unpaired chains have index values of 0. Complete SC data from the Immune peptide Database (IEDB) was also added to our dataset. In total our SC dataset comprised 1,447 TCRs, BS α 21,207 chains, and BS β 25,927 chains.

### CDR information

Our method for predicting the specificity of TCRs requires TCR gene family and complete CDR3 sequences. To obtain this, we retrieved all human germline CDR loop information from the International ImMunoGeneTics Information System Gene database (IMGT/GENE-DB).^10^CDR1 and CDR2 loops can be retrieved directly from the database. However, CDR2.5 needs to be extracted from the IMGT alignment sequence, and is defined by the residues in columns 81-86 of the gapped alignment (F+ORF+in-frame P amino acid sequences with IMGT gaps), as discussed in Dash et al^2^. After translating the data to protein sequence, we produced non-redundant datasets by removing duplicate TCRs (sequence identity < 100%).

### Baseline method

We implemented TCRdist^2^ as a baseline method for classifying TCRs according to their peptide specificity. TCRdist is based on a nearest-neighbor approach, where the distance between TCRs is obtained from protein sequence alignment scores between TCRs. Using a BLOSUM62 matrix, a protein alignment is performed between any two TCRs using CDR loops 1, 2, 2.5, and 3. Subsequently, CDR 1, 2, and 2.5 are given a weight of 1, where CDR3 is given a weight of 3. Finally, the weighted sum of the CDR loop alignment scores is used as a proximity measure, and TCRs are assigned the peptide specificity of their nearest neighbor.^2^

### SwarmTCR

The main idea behind SwarmTCR is that the “importance” of the α and β chains as well as the CDR regions within these chains varies depending upon the peptide that is being recognized, as described in the literature.^7^ In order to reflect this, SwarmTCR learns optimal weights for each of the eight CDR loops in a peptide-specific fashion. SwarmTCR explores the eight-dimensional (with SC data) or four-dimensional (with BS data) space of CDR weights with Particle Swarm Optimization (PSO), an established optimization technique inspired by the natural flocking behavior of birds that has been shown to achieve good performance in a wide range of optimization contexts.^11^

The weights are used in a nearest-neighbor framework as done in TCRdist. We framed this as an optimization problem, where the objective is to identify a set of weights that maximize classification performance as measured by Average Precision (AP) (eq. 1). AP was selected as the objective function to address the issue of unbalanced datasets, as suggested by Saito and Rehmsmeir.^12^ We used Particle Swarm Optimization (PSO) for carrying out the optimization of the weights and maximize AP on a training set.

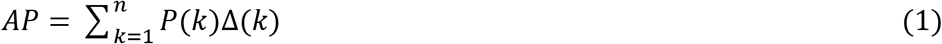

AP is determined by the sum over every position of the precision-recall curve where *k* is the rank of the retrieved TCRs, *n* is the number of TCRs, P(*k*) is the precision at cut-off *k* and **Δ***r*(*k*) is the change in recall from *k - 1* to *k*.^13^

In PSO particles are initially placed in a multidimensional space at random, with each particle representing a possible solution to the optimization problem. At each iteration, particles move with a velocity vector that is a function of both the local best of the particle and the global best. The velocity (**v**) and position (**p**) of a particle *i* are updated at each time step *t* according to eq. 2 and 3:

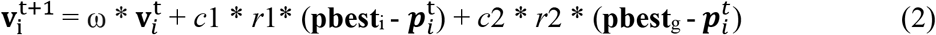

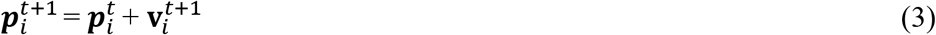

where ω is an inertia factor set to 0.5, *c1* and *c2* are scaling factors set to 0.5 *r*1 and *r*2 are two random numbers between 0 and 1, **pbest**_i_ is the position of particle *i* that has resulted in the best value for the objective function so far, while **pbest**_g_ is the global best (i.e., the position corresponding to the best value so far across all particles).

The optimization is set to terminate if the swarm moves ≤ 10^-8^ from its best position or if the change in the swarm’s best objective value is ≤ 10^-8^. The swarm size is set to 25, with a maximum number of 20 iterations.

Recent literature^7^ has shown that the use of α and β chains is not always equal, therefore we reason that the weighting scheme should not be either. Here we propose the use of PSO to determine the weights applied to the alignment distance values for each class of peptide specific TCRs. Additionally, this method does not assume any preference for CDR looping regions, where weights can be a floating-point number between 0 and 1. In our implementation, the metric chosen for optimization was the average precision for the K-nearest neighbor classifier.

### The SwarmTCR model

We define as “training set’” the TCRs used to obtain an optimal set of weights maximizing average precision, and test set as the TCRs where the performance of the optimal weights is evaluated. Within both sets, we have a reference subset containing labeled TCRs and a sample subset that the nearest neighbor approach compares against the reference subset to infer peptide labels for the TCRs.

Training and test sets for SC and BS data were constructed differently due to data availability, with BS data being much more abundant than SC data. For BS, the training and test sets were filled using a 50/50 split, and for both training and test sets half of the TCRs specific for a particular peptide were placed into the reference subset and the remainder into the sample subset (Figure 2A).

**Figure 2.**
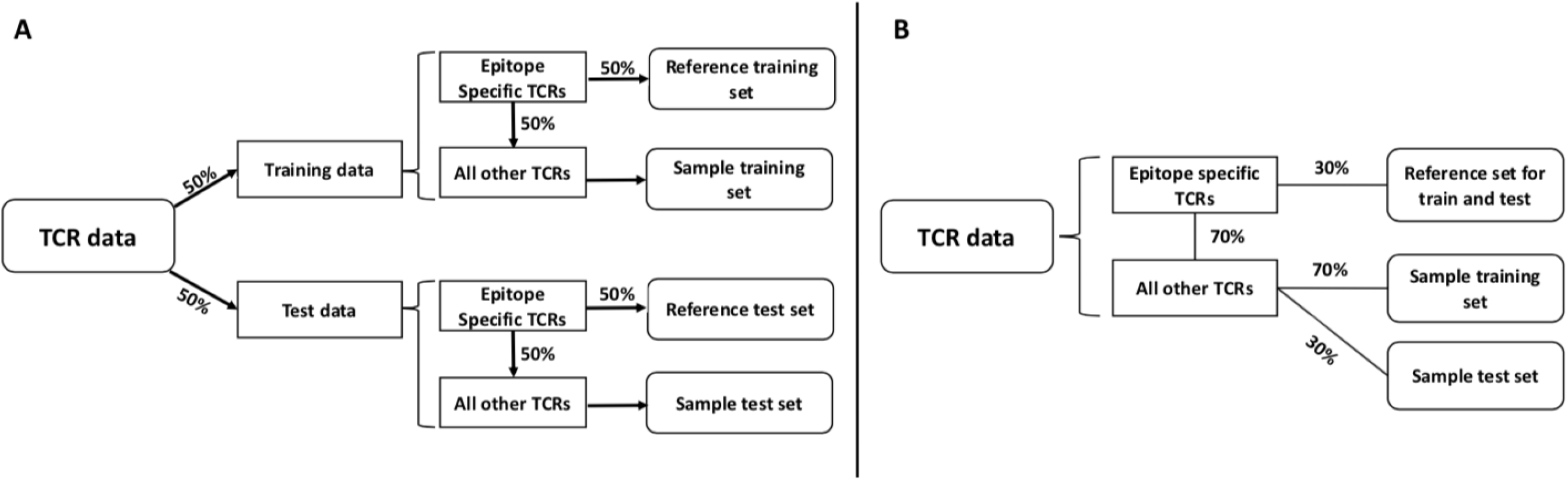
The allocation of data for both BS (A) and SC (B) model is illustrated above. All TCRs are split proportionally into the sample and reference sets. As described in the literature, the same SC reference set is used in training and testing (due to abundance of data, this was not necessary in the BS analysis). The subset labeled peptide specific TCRs are the group of TCRs specific for the peptide tested by SwarmTCR.

For SC, 30% of all TCRs specific for a peptide were placed into the reference subset for both training and testing sets. The same reference subset was used in training and testing due to limited amounts of SC data. In order to create the sample subsets for training and testing, the remaining 70% (TCR specific) undergoes another 70/30 split (Figure 2B). We note that the sample reference sets are distinct for training and test. We also ensured that the different proportions of TCR peptide specificities were equally represented in training and test sets.

Once data was randomly allocated into the training and test sets as described above, we performed the PSO procedure on the training set. Each solution (optimal set of weights maximizing average precision) was then applied to the test set. Cross-validation was performed using repeated random sub-sampling for 50 iterations on both SC and BS datasets.

## Results

### Classification performance of SwarmTCR

Figure 3 shows two crystal structures of TCR/pMHC complexes to visually illustrate the fact that the α and β chains can be involved in pMHC binding to a very different extent, depending on the peptide that is being recognized. In the example shown in Figure 3, one TCR (PDB ID: 4G8G^14^ has 16 (59%) α chain residues and 11 (41%) β chain residues in contact with the pMHC, whereas the other^15^ has 9 (39%) α chain residues and 14 (61%) β chain residues in contact with the pMHC. This is consistent with results in the literature.^7^

**Figure 3.**
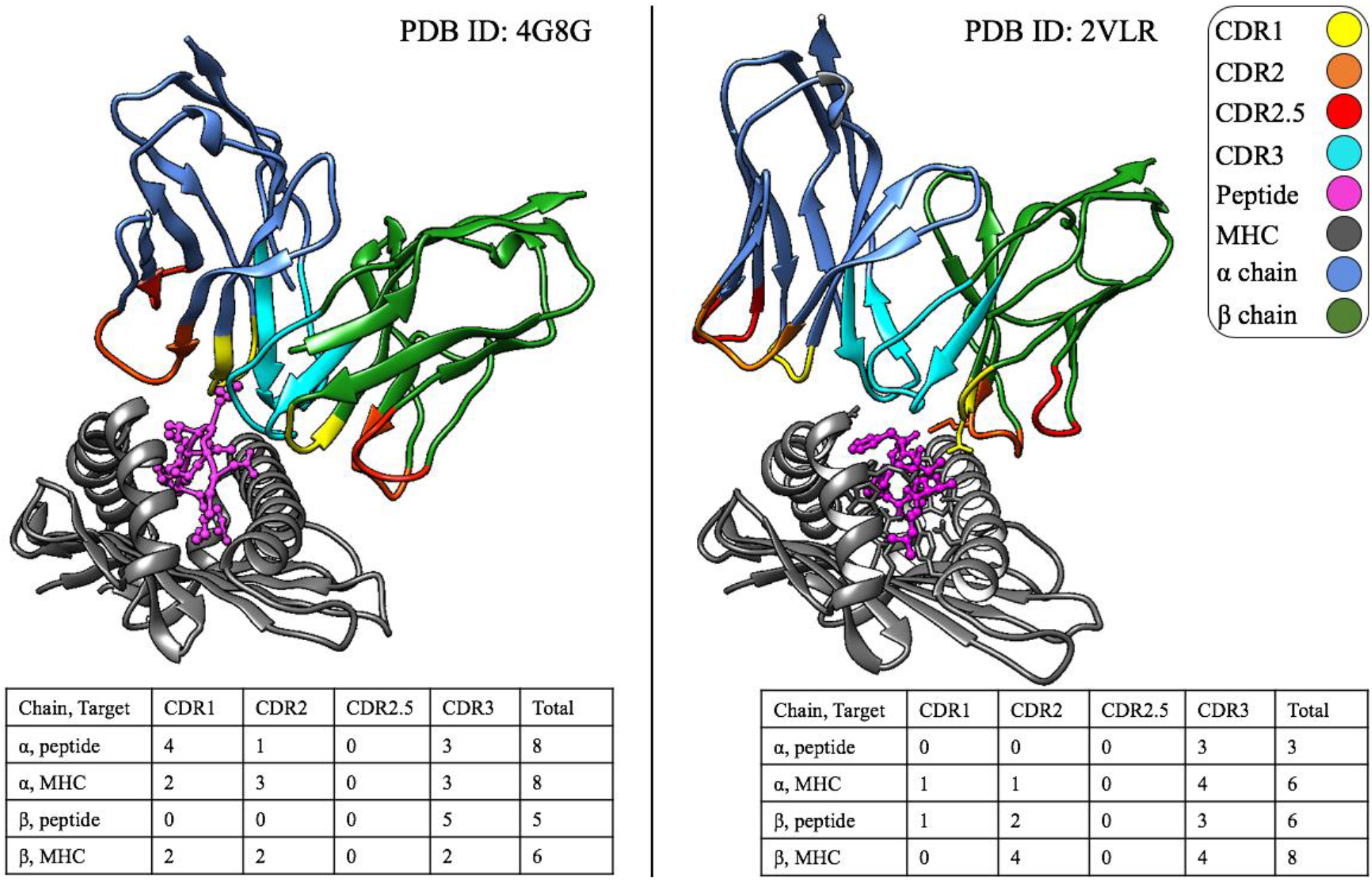
The complexes shown here (PDB ID: 4G8G, peptide: KRWIILGLNK and PDB ID: 2VLR, peptide: GILGFVFTL) express the need for SwarmTCR. The 4G8G complex illustrates an α driven interaction and 2VLR conversely, a β driven interaction. All protein chains (including the CDR loops) are color-coded to supplement the tables beneath each structure. The tables show the number of contact residues in each CDR loop and a target

Based on this observation, SwarmTCR optimizes the weights used to compute the CDR alignment scores underpinning the nearest-neighbor classification approach. In contrast, previous attempts at predicting TCR specificity (TCRdist method) used a static weighting scheme with equal α and β chain contributions and fixed CDR loop weights.^2^ The SwarmTCR method makes no assumptions about chain or CDR loop importance, but learns the weights in a peptide-specific fashion.

Mean and standard deviation of the optimized weights for several peptides are shown in Figure 4, together with classification performance for SwarmTCR and TCRdist^2^ separately for SC and BS data.

**Figure 4.**
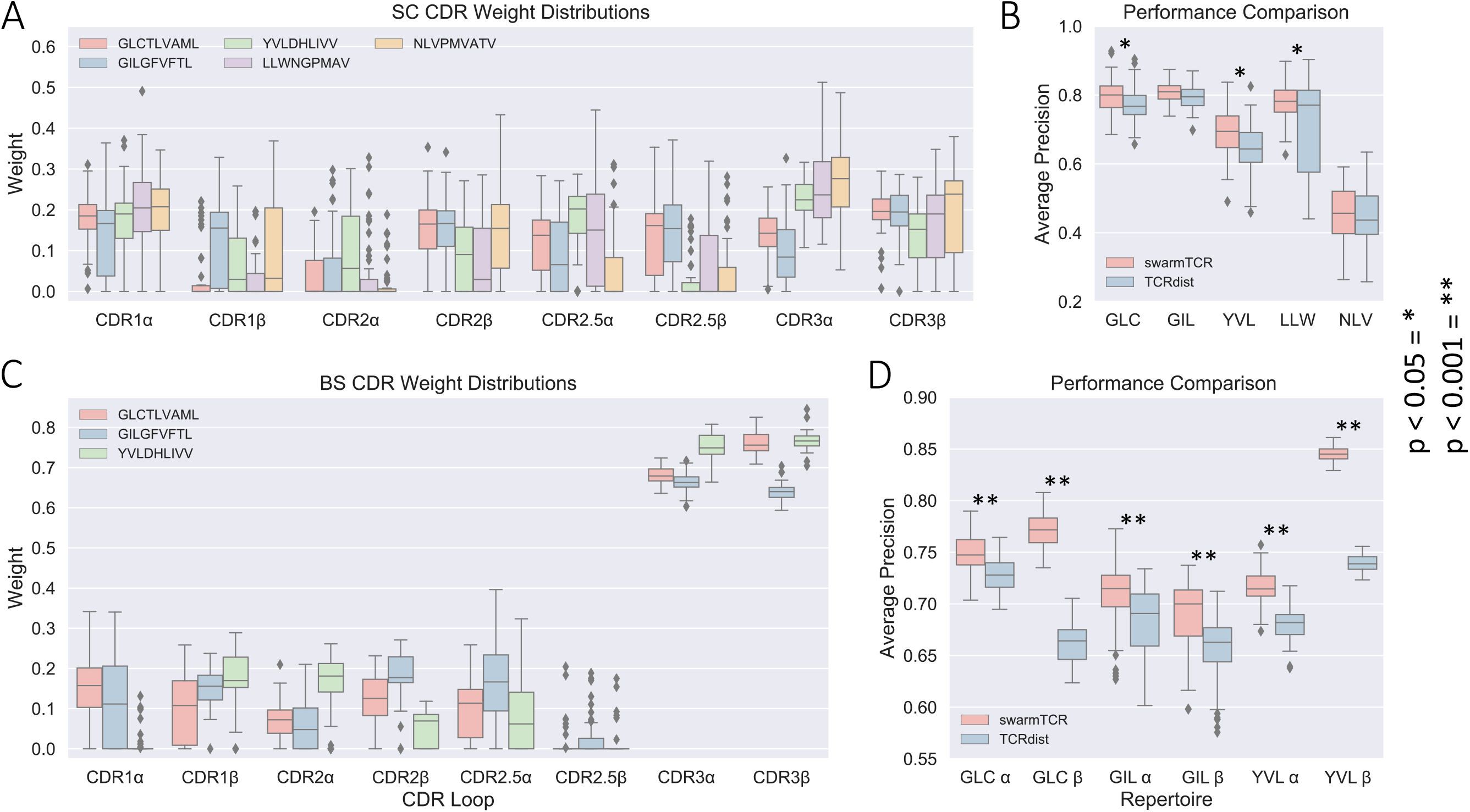
These boxplots summarize the results of SwarmTCR and compare them against TCRdist. (A, C) SC and BS SwarmTCR results describe weights (y-axis) selected for each CDR loop (x-axis) for each repertoire tested. (B, D) SC and BS performance comparison of SwarmTCR and TCRdist compare average precision scores (y-axis) for each repertoire tested (x-axis). P-values for performance comparison are defined by two-sample independent t-test.

### Single cell sequencing

SC data provides paired α/β chain information, i.e., the complete TCR sequence is available. The SwarmTCR optimization procedure for SC data involves the use of eight separate weights, since we have paired α and β chain sequences. The results of our SC analysis show relatively high weight being placed on non-CDR3 loops, although the CDR3 region has high weight for several peptides (Figure 4A). Interestingly, in the case of the EBV YVL peptide and the Yellow Fever LLW (peptide: LLWNGPMAV) peptide the SwarmTCR optimization procedure assigns more weight to the α chain, suggesting that the α and β chains might have a more or less prominent role in TCR peptide recognition depending on the peptide, which is consistent with the example shown in Figure 3 and the previous literature.^7^

By looking at the results in Figure 4B and in Figure 5B, we can see that the largest difference between the classification performance of TCRDist^2^ and SwarmTCR is for the EBV YVL and GLC peptides. The optimized weights for these peptides differ substantially from the fixed TCRdist weights. Based on the optimized weights, YVL appears to favor the α chain as noted above, with only CDR2β being weighted more than its α counterpart.

**Figure 5.**
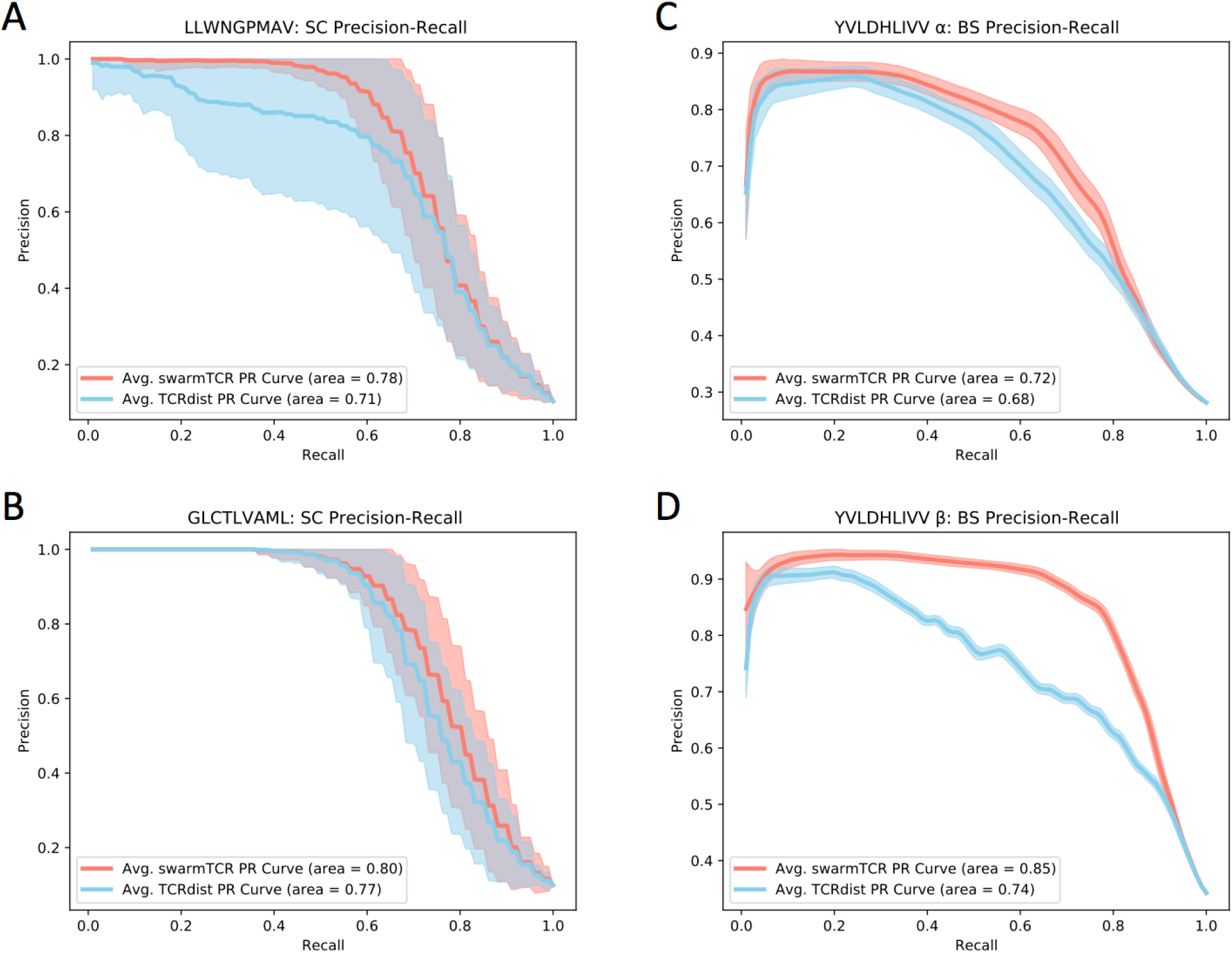
These precision-recall curves show the performance of SwarmTCR and TCRdist on the data used for 50 cross-validation iterations. TCRdist mean curves are in blue and SwarmTCR mean curves are in red, while the shaded regions cover one standard deviation.

### Bulk Sequencing

The SwarmTCR optimization procedure was carried out in the same manner as for SC, except that the weights to optimize are four instead of eight, since we have unpaired α or β chain sequences. Compared to SC data, the results on BS data show more weight being placed on CDR3 loops, indicating its importance in predicting the specificity of TCR data when using only one chain. While TCRdist assigns to the CDR3 loop three times the weight of the CDR1, CDR2, and CDR2.5 regions^2^, SwarmTCR assigns to CDR3 between ~4 and ~64 times the weight of the other regions (using the average weight as a measure), as it can be seen in Figure 4C. These differences between the SwarmTCR weights and the original weights by Dash et al.^2^ have a substantial impact on the classification performance for the GLC and YVL peptides using the β chains (Figure 4D).

Figure 5 (C and D) shows precision-recall (PR) curves for a representative peptide (EBV YVL), obtained by averaging 50 curves, with the shaded region representing one standard deviation above and below the mean. SwarmTCR outperforms the original weights used in TCRdist^2^ for both chains, with a more substantial improvement for the β chain (AUCPR 0.85 with the optimized weights vs. 0.74 with the original TCRdist weights).

## Discussion and Conclusion

We introduced SwarmTCR, a computational approach for predicting TCR specificity that maximizes classification performance within a nearest neighbor framework by identifying optimal CDR weights. Compared to the results obtained with fixed TCRdist weights, SwarmTCR performs consistently better, with some peptides showing more substantial improvement than others (Figure 5). We note that in a worst-case scenario SwarmTCR can always fall back on the weights used by TCRdist if those yield maximum performance during the PSO step.

When comparing CDR weights in SC and BS data, we noticed stark differences between the results obtained with the two data types. In particular, we found that SwarmTCR assigns much more weight to the CDR3 region in BS data, whereas the results on SC data show relatively higher weights for the germline CDR loops. Due to the small size of the SC dataset, the diversity of TCR gene families is likely considerably lower than that found in the BS dataset. Therefore, the lower gene family diversity in the SC dataset compared to BS could partly explain the higher predictive power of gene family (germline loops) in SC data. Another reason for this difference in the weights between the two data types is the presence of paired chains information in SC, where combinations of TCR genes for α and β chains would likely be selected for by the optimization approach. More SC data is needed to further elucidate the issue. Consistent with the substantial differences in size between the two datasets, SC results show higher variance in both performance and weight selection than BS results.

An important question to consider is whether the optimized weights can also be interpreted to reflect chain and CDR usage. In other words, if a chain or a CDR region receives a high weight during the optimization step, does that mean that it also makes a large number of contacts with the pMHC? Our results suggest that the optimized weights can point to possible TCR chain and CDR loop usage, as shown in Figure 4 A and Supplementary Table 2 for the GIL TCR/pMHC crystal structure (Figure 3, PDB ID: 2VLR) with respect to β chain dominance and CDR2β loop usage (Table 1).

**Table 1.** This table compares SwarmTCR’s mean weights from our SC GIL analysis and actual contacts from Figure 3 PDB ID: 2VLR. Columns 2-5 detail CDR loops for α (rows 2 and 3) and β (rows 4 and 5).

| CDR Loop | CDR1 | CDR2 | CDR2.5 | CDR3 |
| --- | --- | --- | --- | --- |
| Structure Contacts $\alpha$ | 1 | 1 | 0 | 7 |
| SwarmTCR Mean Weight $\alpha$ | 0.140 | 0.057 | 0.093 | 0.095 |
| Structure Contacts $\beta$ | 1 | 6 | 0 | 7 |
| SwarmTCR Mean Weight $\beta$ | 0.121 | 0.156 | 0.145 | 0.191 |

A recent study^16^ corroborates these findings, explaining CDR1β and CDR2β’s role in pMHC recognition, as well as CDR3β’s conserved arginine fitting into a pocket between the peptide and the MHC α2-helix. Additionally, this study explains CDR3β’s sequence conservation and notable variability in CDR3α. This likely explains the weighting results of SwarmTCR (GIL, Figure 4 and Table 1) despite the number of contacts in the 2VLR TCR/pMHC structure. We also note that SwarmTCR’s weight results for YVL and LLW repertoires align with findings from this study, indicating the importance of the α chain in pMHC recognition.^16^

However, one has to exercise caution when interpreting the weights in a structural sense. As discussed above, the weights are the result of an optimization process designed to maximize classification performance, and factors other than structural importance can play a role in determining the optimal weights. If we consider the crystal structures and literature mentioned, this is shown by our BS weighting results and differences present in Table 1. Nonetheless, given large amounts of TCR sequence data, peptide-specific optimal weights can provide helpful information in elucidating TCR/pMHC interactions.

Sequence-based approaches to infer TCR specificity are appealing due to their computational efficiency and the availability of sequence data^2,5,17^ However, structural data continue to provide information that expands and sometimes challenges our current understanding of TCR/pMHC interactions. For example, one study found a strong negative correlation between mean CDR3 α, β charge and peptide charge.^2^ Another study^3^ showed how cross-reactive peptides share similar pMHC features (structural motifs and electrostatic potential) despite having different peptide sequences. These findings point to the importance of factoring in structural information for further improving prediction methods. However, more work needs to be done both at the experimental level (generation of more crystal structures) and the computational level (reliable and scalable modeling of TCRs and pMHC complexes).

Being able to reliably predict TCR specificity will push the boundaries of many disciplines including vaccine design, immunotherapy, cancer research, and disease detection/prevention in new directions. Here we have introduced SwarmTCR, a nearest-neighbor approach that optimizes CDR weights by maximizing classification performance. SwarmTCR was benchmarked on both SC and BS data, and compared against the most recent state-of-the-art methodology, TCRdist. The results showed that SwarmTCR performs better on both SC and BS datasets.

## Acknowledgements

We thank Robin Brody for technical assistance and Dr. Matteo Ligorio and Sean West for reading the manuscript.

